# Divide and Conquer: Real-time maximum likelihood fitting of multiple emitters for super-resolution localization microscopy

**DOI:** 10.1101/659631

**Authors:** Luchang Li, Bo Xin, Weibing Kuang, Zhiwei Zhou, Zhen-Li Huang

## Abstract

Multi-emitter localization has great potential for maximizing the imaging speed of super-resolution localization microscopy. However, the slow image analysis speed of reported multi-emitter localization algorithms limits their usage in mostly off-line image processing with small image size. Here we adopt the well-known divide and conquer strategy in computer science and present a fitting-based method called QC-STORM for fast multi-emitter localization. Using simulated and experimental data, we verify that QC-STORM is capable of providing real-time full image processing on raw images with 100 µm × 100 µm field of view and 10 ms exposure time, with comparable spatial resolution as the popular fitting-based ThunderSTORM and the up-to-date non-iterative WindSTORM. This study pushes the development and practical use of super-resolution localization microscopy in high-throughput or high-content imaging of cell-to-cell differences or discovering rare events in a large cell population.

## 1. Introduction

Super-resolution localization microscopy (SRLM) provides a remarkable improvement in the spatial resolution of far-field fluorescence microscopy with a relative simple setup and user-friendly experimental procedures, thus becomes an important tool for various biomedical researches [1]. The resolution improvement of SRLM is usually achieved upon sparse excitation and precise localization of a large number of fluorescent emitters. Therefore, thousands and even tens of thousands of raw images are necessary for obtaining a full list of molecule localizations that is further used for reconstructing a final super-resolution image with a sufficient sample frequency [2-4].

It is widely accepted that the imaging speed and/or imaging throughput of SRLM can be significantly improved by allowing multiple emitters, rather than only one emitter, to be localized within a region-of-interest (ROI). However, the reported multi-emitter localization algorithms normally run at a very slow speed, typically several hours for a super-resolution image with a small field of view (FOV) [5]. Recently, Ma et al presented a non-iterative multi-emitter localization algorithm, called WindSTORM [6], which runs at a very fast speed. Actually, WindSTORM is capable of localizing experimentally up to 0.7 million emitters per second for bright emitters without compromising localization accuracy, thus enabling real-time image processing for an FOV of 40 µm × 40 µm with a spatial resolution of 53 nm (measured by the width of the microtubules in a reconstructed image). However, WindSTORM is still not sufficient to explore fully the fast imaging power of popular sCMOS cameras used in SRLM, because these fast cameras can support an FOV of up to 200 µm × 200 µm (when using a typical pixel size of 100 nm) and an exposure time of 10 ms [7]. And, currently WindSTORM is not capable to process 3D raw images.

Instead of producing a super-resolution image from a full list of molecule localizations, researchers are now exploring the use of deep learning to provide a super-resolution image directly from raw images or from a much smaller number of molecule localizations [8]. For example, Deep-STORM has been developed to use a deep convolutional neural network to create a super-resolved image from raw images directly, and thus achieves up to three orders of magnitude faster speed than the conventional localization-based methods [9]. However, the size of the neural network and the available memory of commercial graphics processing unit (GPU) limit the maximum size of raw images that can be processed by Deep-STORM. Another deep learning based method, called ANNA-PALM, uses machine learning to generate a super-resolution image from sparse localizations or even diffraction-limited input images, and thus massively accelerates the imaging speed [10]. However, ANNA-PALM requires a long post-processing time. And, it is reported that the performance of ANNA-PALM is affected by the amount and variety of training data, and that an effective algorithm is necessary to be used to identify and reduce image artifacts [10]. For some applications where the training data is not easy to be obtained or even unavailable (for example, rare clinical samples with unknown structures), the traditional localization-based approaches would be more suitable and/or more reliable due to the lower risk of image artifacts.

Taking the above discussions into consideration, we conclude that exploring new multi-emitter localization algorithm for enhancing the imaging speed without compromising the spatial resolution and the user-friendly features of SRLM is still on heavy demand. The image analysis speed is preferable to be real-time, so that the acquisition parameters can be optimized during SRLM experiments [11-13]. Note that high imaging throughput and real-time acquisition optimization are both required for applying SRLM in high-throughput or high-content imaging of a large cell population [5, 11, 14, 15], which are currently used to reduce selection bias, reveal cell-to-cell differences, and discover rare phenotypes or events [16, 17]. Of course, it would be interesting to investigate how to combine localization algorithms with deep learning to further improve the imaging speed and/or imaging throughput of SRLM, without compromising other requirements from high-throughput localization microscopy.

In this paper, we propose a new method, called QC-STORM, for fast maximum likelihood estimation (MLE) of the locations of multiple emitters. Using the well-known divide and conquer strategy in computer science, QC-STORM firstly extracts ROIs from raw images, and then sends them to a series of MLE-based localization algorithms that are designed to provide fast image processing on ROIs containing a fixed number of emitters. Using simulated and experimental datasets, we compare the localization performance among QC-STORM, ThunderSTORM and WindSTORM. We verify experimentally that QC-STORM enables real-time data processing for an FOV of 100 µm × 100 µm and an exposure time of 10 ms, with similar or even better spatial resolution than that from the popular fitting-based ThunderSTORM and the non-iterative WindSTORM. We further combine QC-STORM with ANNA-PALM to triple the imaging speed. The open-source codes and the QC-STORM plugin for ImageJ and Micro-Manager are available at: https://github.com/SRMLabHUST/QC-STORM.

## 2. Methods

### 2.1 MLE_bfgs_ for fast localization of sparse emitters

Among the numerous reported molecule localization algorithms, it is widely accepted that the MLE-based algorithms are able to achieve the highest localization precision [18]. However, MLE-based algorithms usually suffer from high computational intensity [18], and thus are normally combined with GPU computation for a faster localization speed. Here we make several optimizations to the mathematical model and the GPU computation of a conventional MLE-based algorithm, so that the localization speed of this algorithm can be further improved. Similar to previous works [19, 20], we use Gaussian function to model the point spread function (PSF) of our imaging system. For astigmatism 3D imaging, the observed signal model is:

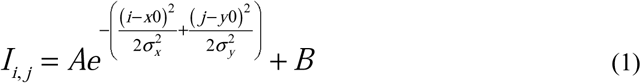

here, *I*_*i,j*_ is the theoretical signal intensity at pixel (*i, j*), *A* and *B* are the peak signal and background intensities, respectively, (*x*_*0*_, *y*_*0*_) is the center position of the fluorescent molecule, and σ*_x_* and σ*_y_* are standard deviation width of the Gaussian PSF along horizontal and vertical directions, respectively. For 2D PSF model, the two widths are merged into a single width.

Practically, since the observed signal is usually contaminated by noises (mainly photon shot noise), based on maximizing the probability of the observed signal [20], the MLE optimization problem for finding the molecule center by minimizing the loss is derived as:

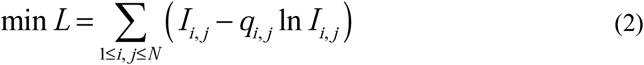

here, the *q*_*ij*_ is the observed signal intensity at pixel (*i, j*), and *N* is the size of the extracted ROI. Taking the fluorescence signal intensity into account, the optimization target *L* is a negative number with a big absolute value (typically larger than 10^5^). If *L* is represented by a single-precision floating-point number, the accuracy of the decimal part of *L* is less than 10^−2^, meaning that it is impossible to achieve high accurate gradient for the parameter optimization. Therefore, *L* must be represented by a double-precision floating-point number, which results in an order of magnitude decrease in the computation speed than that using a single-precision floating-point number in typical graphics cards [21].

To increase the computation speed, we apply a key modification to this MLE optimization model:

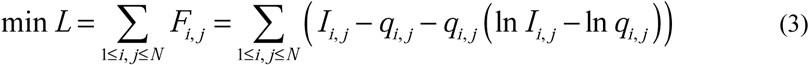

where *F*_*ij*_ is used for simplicity. By subtracting a constant that is calculated from the observed signal, the absolute value of *L* can be theoretically reduced to zero, thus enabling an accurate parameter optimization using a single-precision floating-point number. Moreover, this treatment does not influence the gradient calculation and the following optimization.

Then, we rewrite the PSF model as:

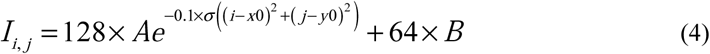

The coefficients in Eq. (4) are empirical scaling factors for ensuring that all fitting parameters are within a similar dynamic range, and thus improving the algorithm robustness [22]. Additionally, the division operation in Eq. (1) is replaced by the multiplication shown in Eq. (4) to accelerate computation.

Finally, the Broyden–Fletcher–Goldfarb–Shanno (BFGS) algorithm [22] is introduced to solve Eq. (3). Therefore, this modified MLE-based algorithm is called MLE_bfgs_. Moreover, the algorithm is accelerated by GPU computation that simultaneous fits thousands of independent molecules. The source codes of implementing the MLE_bfgs_ algorithm are described in: https://github.com/SRMLabHUST/QC-STORM.

### 2.2 Weighted MLE for sparse emitters with signal contamination

The most important task in SRLM is to localize precisely the center positions of the molecules in an extracted ROI. Due to the random activation nature of molecules, the extracted ROI may contain incomplete fluorescence emission signal from adjacent molecules (here we called signal contamination). If the PSF model for molecule localization considers only one molecule, the molecule center cannot be localized precisely (Fig. 1a). In the past, this problem is mainly solved by using a multi-emitter localization algorithm [23]. However, most reported multi-emitter localization algorithms require usually many fitting parameters and a large iteration number, and thus run at a much slower speed than the molecule localization algorithms for sparse emitters.

**Fig. 1.**
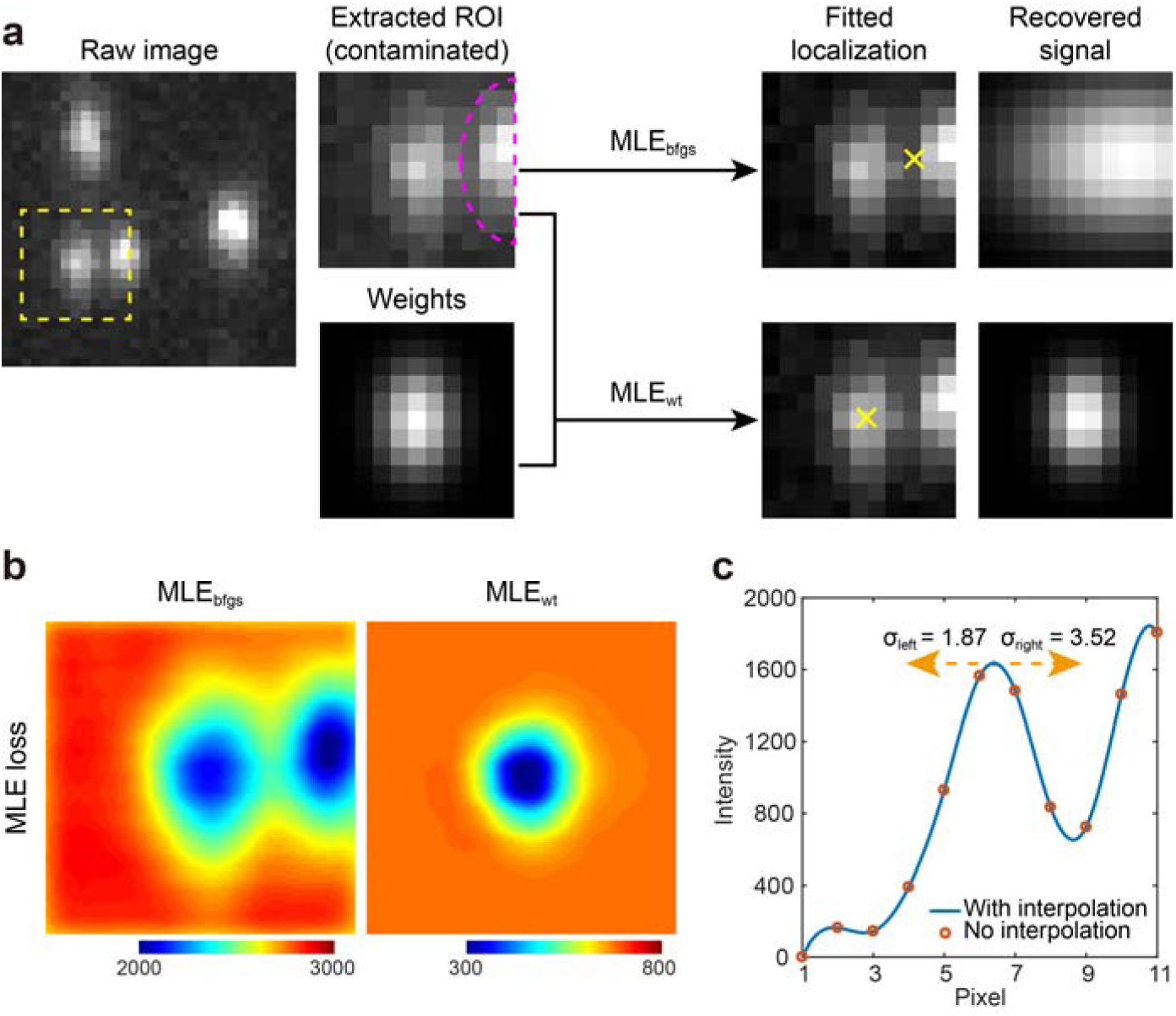
Principle of MLE_wt_ for sparse emitter localization with signal contamination. (a) Comparison on the recovered signal from conventional MLE (MLE_bfgs_, upper) and weighted MLE (MLE_wt_, lower). A contaminated ROI (11 × 11 pixels) is extracted from a raw image taken from astigmatism 3D imaging, and is processed using either MLE_bfgs_ or MLE_wt_. The signal is recovered using the fitted parameters. Note that this ROI can be also fitted using a multi-emitter model at the expense of a slow data processing speed. (b) MLE loss maps from an ROI with signal contamination. In both maps, the fitting parameters other than x and y positions were fixed and adopted from the fitted result of MLE_wt_. (c) Averaged horizontal intensity profile of the extracted ROI in (a). The σ_left_ and σ_right_ denote the estimated standard deviation width (in pixel) for the Gaussian weight function in the left and right sides, respectively.

Weighted maximum likelihood estimation (WLE), which applies a larger weight for high reliable data and a lower weight for low reliable data, is an efficient method to suppress the signal contamination. Previously, WLE has been used to increase the robustness of MLE-based fitting and to reduce bias results from contaminated signal [24, 25]. In SRLM, WLE has also been used for improving localization precision by correcting the PSF distortion in 3D imaging [26]. Typically, different weights should be developed according to the types of application and contamination.

Here we introduce a Gaussian function weighted MLE (MLE_wt_) to suppress the signal contamination from adjacent molecules. As illustrated in Fig. 1a, for a target molecule that doesn’t overlap severely with an adjacent molecule, the use of a traditional MLE-based algorithm presents apparently incorrect localization results. In contrast, if we first estimate the PSF width of the target molecule, and then apply a weight function, which is Gaussian function and is based on the estimated PSF width, to the MLE-based algorithm for sparse emitters, the localization precision can be significantly improved (Fig. 1a). From the MLE loss maps (Fig. 1b), we found that there are two local minima for MLE_bfgs_, but only one for MLE_wt_. Since the goal of MLE is to find the best parameters by minimizing the loss (Eq. (3)), the two local minima (as shown in Fig. 1b) prevent the use of MLE_bfgs_ to find the correct center position of the target molecule.

Below is a brief description of MLE_wt_. Firstly, we apply a Gaussian weight function to the MLE optimization model (see Eq. (3)):

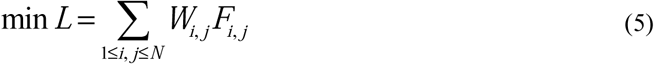

The Gaussian weight function, *W_i, j_*, is calculated using:

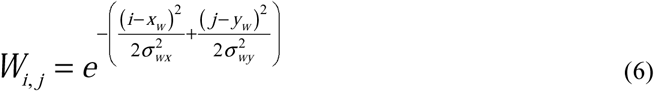

where (x_w_, y_w_) is the center position of the Gaussian weight function, and is directly assigned by the center of the extracted ROI since the target molecule is made to be centered in this ROI during the ROI identification and extraction step. σ_wx_ and σ_wy_ are the estimated widths of the PSF in horizontal and vertical directions, respectively. For 2D imaging, the smaller one of them is considered as the PSF width.

To estimate the PSF width of the target molecule, we first average the signal of the ROI along horizontal and vertical directions, respectively, to obtain two one-dimensional Gaussian shape curves (Fig. 1c). Then, the standard deviation widths of the Gaussian shape curves in four sides (including left, right, top and bottom) are calculated using the relationship among Gaussian integral, the area and the peak of the Gaussian shape curves:

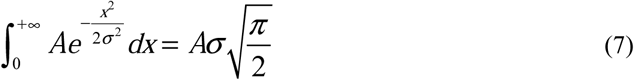

To better calculate the area of the Gaussian shape curves, we apply a 10-fold cubic spline interpolation before the numerical integration of the Gaussian shape curves. Finally, the smaller value of the estimated widths in the left and the right sides is used as the final horizontal PSF width, and the smaller value of the estimated widths in the top and the bottom sides is used as the final vertical PSF width.

An extracted ROI is determined to be contaminated by adjacent molecules if the estimated width of the corresponding Gaussian shape curves in one side is 40% larger than that in the other side. For contaminated ROIs, the estimated PSF widths of the Gaussian shape curves divided by 1.2 are used as the widths of the final Gaussian weight function. For uncontaminated ROIs, the same unit weights are used for all pixels. These parameters are empirically determined to avoid incorrect classification on the uncontaminated molecules. And, since the center of the weight function is not absolutely matched with the ground-truth position of the target molecule, applying MLE_wt_ to uncontaminated molecules may reduce localization precision. Moreover, for truly contaminated molecules, a narrower weight function is helpful for improving localization precision, which is determined by the distance from nearest neighbor molecules.

Finally, Eq. (5) is solved using the same optimization method as that for Eq. (3). Because the Gaussian weight function is calculated before the iterative optimization process for solving Eq. (5), MLE_wt_ runs at a similar speed as the unweighted MLE-based algorithm, MLE_bfgs_.

### 2.3 MLE for two or three emitters

The MLE-based algorithm for sparse emitter localization can be extended to analyze ROIs with multiple emitters if we add multiple PSFs to the observed signal model [27]. For astigmatism 3D imaging, the signal model is:

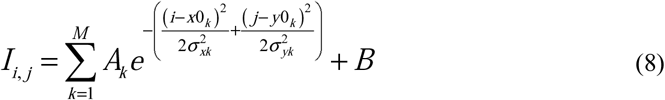

here the *M* is the maximum number of fitted molecules. For 2D imaging, the two widths are merged into a single width. The main difference between multi-emitter MLE and sparse-emitter MLE is that the former has more fitting parameters, and thus requires a larger iteration number. Practically, the iteration numbers and the fitting parameters used in this study are: 5 parameters and 8 iterations for 2D sparse localization, 6 parameters and 11 iterations for 3D sparse localization, 8 parameters and 13 iterations for 2D two-emitter localization, and 11 parameters and 16 iterations both 2D three-emitter localization and astigmatism 3D two-emitter localization, respectively. The MLE-based algorithms for two-or three-emitter localization are solved using the same optimization method as that for MLE_bfgs_.

### 2.4 ROI identification and extraction

The ROI identification and extraction is modified from a reported algorithm called MaLiang [20]. The modifications mainly include: 1) A full GPU computation is used to avoid frequent memory copy between CPU and GPU; 2) A larger annular filter (9 × 9) is used for non-uniform background removal, which is necessary for astigmatism 3D imaging; and 3) A different method is developed to determine the threshold for ROI identification in raw images with high emitter density, since directly calculating the standard deviation of background-removed images will result in overestimation of background noise intensity. The workflow is shown in Fig. 2. For the last modification, we equally split the background-removed image into four small images with the same size, and then generate a noise intensity image where the pixel values equal to the smallest pixel values in the same positions of the four images. This treatment ensures rejection of fluorescent signal from this raw image because of the random activation nature of fluorescent emitters during experiments (that is, it is almost impossible to have emitters at the same position of all four images). The threshold for ROI identification is calculated by 8.5 times the standard deviation of the noise intensity image. After the smoothed image is enhanced by a 3 × 3 high-pass filter, where the center pixel is set to be 2 and the other pixels are set to be −0.125, the targeted molecules are identified based on this threshold and local maximum. The high-pass filter is not used for processing astigmatism 3D raw images. Finally, ROIs of the targeted molecules are extracted from the raw images. In this study, the ROI sizes are set to be 7 × 7 for 2D, and 11 × 11 for astigmatism 3D raw images, respectively.

**Fig. 2.**
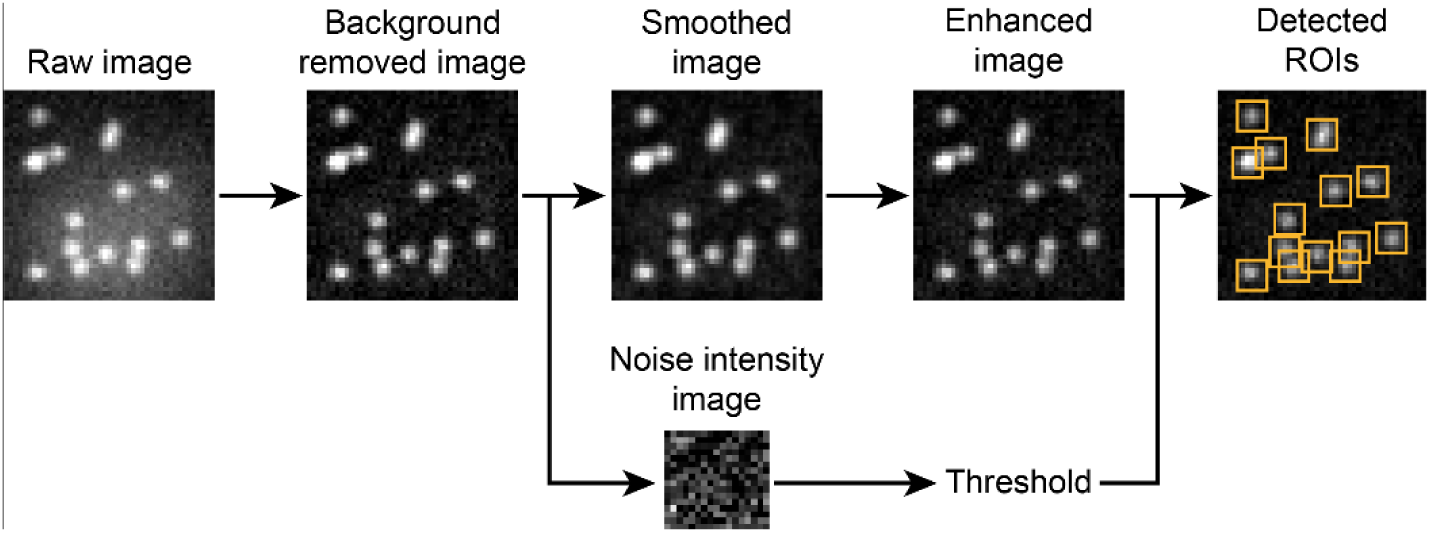
The workflow for ROI identification and extraction.

### 2.5 QC-STORM plug-in

We developed a Java-based user-friendly plug-in, called QC-STORM, to include all the image processing steps for fast MLE fitting of sparse and multiple emitters, as well as super-resolution image reconstruction and some image statistics. The image reconstruction is based on the popular Gaussian rendering method, where the width of Gaussian kernel is determined by localization precision. This plug-in can be easily integrated into the widely-used ImageJ software for off-line image processing or the Micro-Manager software for online image processing. Note that the performance of QC-STORM mainly depends on the floating-point operation speed of GPU and the memory bandwidth of CPU. By combing a Dell precision T7610 workstation with a good graphics card (see Section 2.7 for details), QC-STORM is capable of providing real-time processing for raw images with a size of 1024 × 1024 pixels and an acquisition time of 10 ms. QC-STORM is also capable of providing statistical information, including mainly emitter brightness, background, signal-noise-ratio (SNR), localization precision and PSF widths, for real-time acquisition optimization during SRLM experiments.

### 2.6 Sample preparation and super-resolution fluorescence imaging

SRLM experiments were performed on an Olympus IX73 inverted optical microscope. A 640 nm and a 405 nm laser were combined and coupled into a multimode optical fiber. The fiber output was collimated and focused onto the biological sample using an oil-immersion objective (100×, NA 1.4, Olympus) to provide a homogeneous flat-field illumination [7]. Alexa 647 or CF 680 labeled U-2 OS cells were imaged with the 640 nm laser at an illumination intensity of 7 kW/cm^2^ while soaked with standard STORM buffer (50 mM Tris, pH 8.0, 10mM NaCl, 10% glucose, 100 mM mercaptoethylamine, 500 µg/mL glucose oxidase, 40 µg/mL catalase) [28]. Images were acquired with a popular sCMOS camera (Flash 4.0 V3, Hamamatsu Photonics) at an exposure time of 10 ms and a pixel size of 108 nm. The FOV is 110 μm × 110 μm.

### 2.7 Image simulation and algorithm evaluation

We simulated 2D and astigmatism 3D raw images with various activation density for algorithm evaluation. For 2D images, Gaussian PSF with a full width at half maximum (FWHM) of 300 nm was used. For astigmatism 3D images, the molecules were randomly distributed within a depth range of 600 nm, and the PSF width in different depths were calculated from the open dataset of real astigmatism 3D microtubule images [29]. For both 2D and astigmatism 3D images, the total photon was simulated to follow a log-normal distribution with a mean of 5000 photons and a standard deviation of 2000 photons, and the background was set as 100 photon per pixel [30]. At each activation density, we simulated 200 images with 256 × 256 pixels and 100 nm pixel size. The molecules were randomly distributed in the images and the number of molecules were calculated by multiplying the image area by the activation density. Moreover, the simulated images considered shot noise and camera specifications (0.77 quantum efficiency, 1.6 e-read noise).

We compared the localization performance of our QC-STORM with ThunderSTORM [27] and WindSTORM [6] using the same simulated image datasets. ThunderSTORM is a widely-used and comprehensive ImageJ plug-in for SRLM, while WindSTORM is a newly developed non-iterative algorithm for fast localization of high-density emitters. ThunderSTORM provides fitting modes for sparse emitter and multiple emitters, respectively. We calculated the Fourier ring correlation (FRC) resolution [31] for experimental raw images. Before calculating the FRC resolution, localizations of molecule that emits fluorescence in consecutive frames were detected and merged into a single localization using a distance threshold of 50 nm. All evaluations were performed on a Dell precision T7610 workstation, which has two Intel Xeon E5-2630 V2 CPU working at 2.6 GHz and 56 GB memory, and a Gigabyte GeForce RTX 2080 Ti Gaming OC graphics card with 11 GB memory.

## 3. Results and discussion

### 3.1 The Divide and Conquer strategy

After ROI identification and extraction using the method described in Section 2.4, we apply the divide and conquer strategy to classify and process the ROIs. The classification is based on two parameters: the distance between the nearest neighboring ROI centers, and the estimated PSF widths. As seen in Fig. 3a, we first need to determine whether an ROI contains only one emitter. For this purpose, we consider two criteria: 1) Whether the distance between the centers of this ROI and its nearest neighboring ROI is larger than a threshold (that is, ROI width / 2 + 1, or 4.5 pixels for the 2D images in this study); and 2) Whether the averaged PSF width estimated by MLE_wt_ along the four directions (see Section 2.2 for details) is smaller than a threshold (that is, ROI width / 2 / 2.35, or 1.49 pixels for the 2D images in this study). The ROI is considered to contain only one emitter if both answers are YES. Otherwise, the ROI is classified to have multiple emitters.

**Fig. 3.**
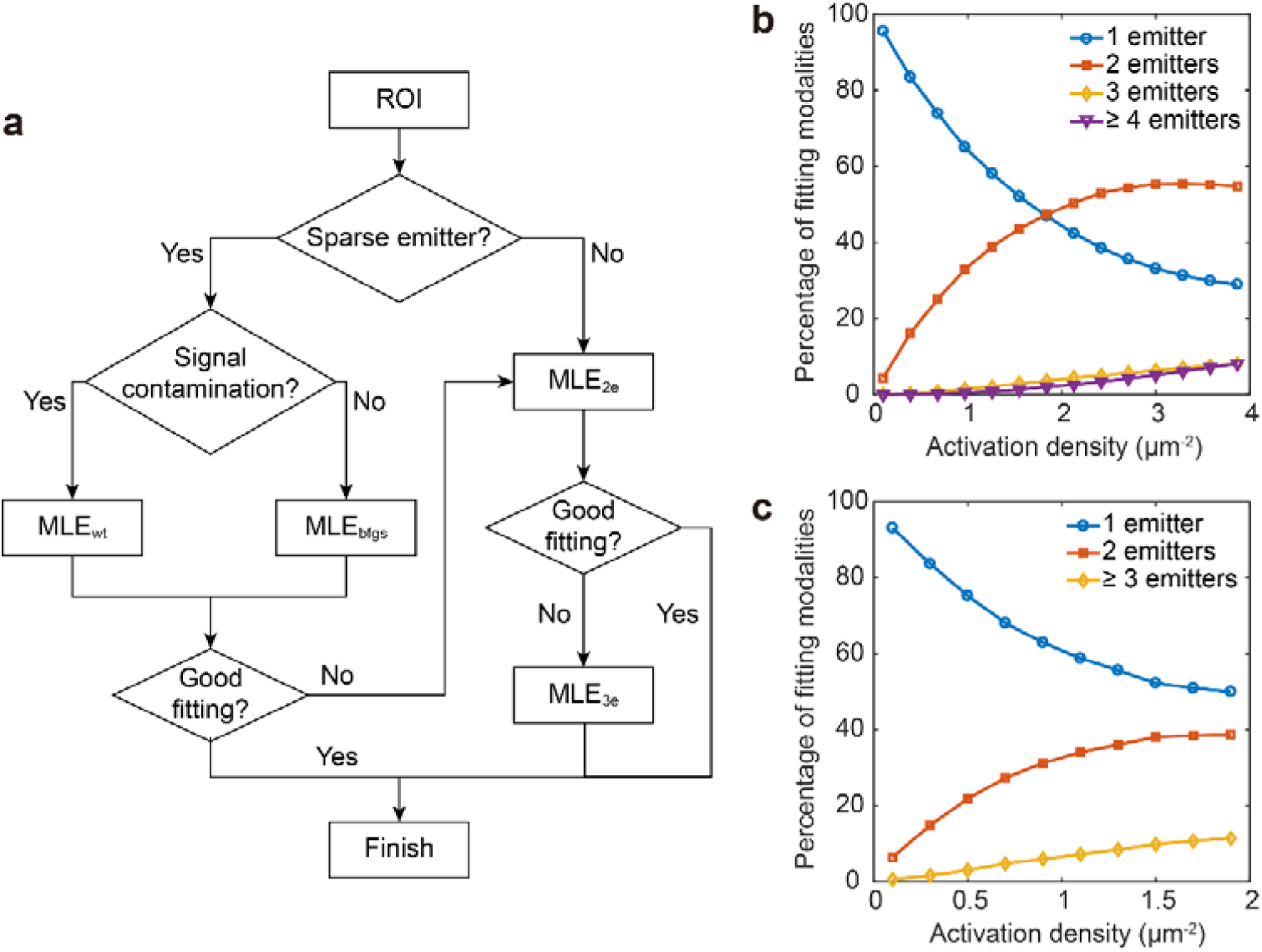
MLE fitting using the divide and conquer strategy. (a) The workflow. (b-c) The percentage of ROIs containing different number of emitters under different activation density. Here (b) is for 2D, and (c) is for astigmatism 3D raw images.

For the ROI considered to have only one emitter, a further judgment related signal contamination is taken, using the criterion described in Section 2.2. Then, the ROI is processed with either MLE_wt_ or MLE_bfgs_, depending on whether the signal is contaminated or not. Since signal contamination commonly results in a larger estimated PSF width, a third judgement related to the fitting quality is taken. If the estimated PSF width is larger than a threshold (that is, ROI width / 2 / 2.35, or 1.49 pixels for the 2D images in this study), the fitting is not considered as a good fitting, and the ROI will be further processed using the two-emitter and three-emitter localization algorithms. The same criterion is used for determining the quality of two-emitter fitting.

To reduce program complexity and ensure real-time image processing under our current hardware configuration, the MLE-based ROI processing in this study is terminated after three-emitter fitting for 2D images, and two-emitter fitting for astigmatism 3D images, respectively. Of course, the divide and conquer strategy shown in Fig. 3a could be extended to include more emitters for higher activation density, if there are more powerful computation resources. Currently, basing on simulated raw images, we found out that real-time image processing could be made possible for raw images with a size of 1024 × 1024 pixels and an acquisition time of 10 ms. In this case, the activation density could be up to 3.5 molecule/µm^2^ for 2D and 1.5 molecule/µm^2^ for astigmatism 3D raw images, respectively, while the rejection rate of ROIs is still less than 10%.

### 3.2 Localization of sparse emitters using simulated datasets

We compared the localization performance among our MLE-based localization algorithms (MLE_bfgs_ and MLE_wt_) and the popular fitting-based ThunderSTORM using simulated datasets. According to Sage et al [32], we used root-mean-square error (RMSE) to quantify the localization accuracy, and Jaccard index to quantify the detection rate. We performed Jaccard index calculations based on the pairing between the found localizations and the ground truth with 100 nm threshold. For 2D simulated datasets, MLE_bfgs_ achieves similar and/or slightly better localization precision (Fig. 4a) and detection rate (Fig. 4b) than ThunderSTORM working at the sparse emitter mode, while a combination of MLE_wt_ with MLE_bfgs_ significantly improves the localization precision and the detection rate in all activation densities. For astigmatism 3D simulation, the superior localization performance of the combination of MLE_wt_ with MLE_bfgs_ over ThunderSTORM is also observed (Fig. 5a-5c). And, it is worthy to point out that, for both 2D and astigmatism 3D simulations, the combination of MLE_wt_ with MLE_bfgs_ provides a doubled maximum activation density than ThunderSTORM, when we use 0.5 as the threshold of Jaccard index. These findings prove the usefulness of the weighted MLE localization algorithm (MLE_wt_) for processing raw images with contaminated signal.

**Fig. 4.**
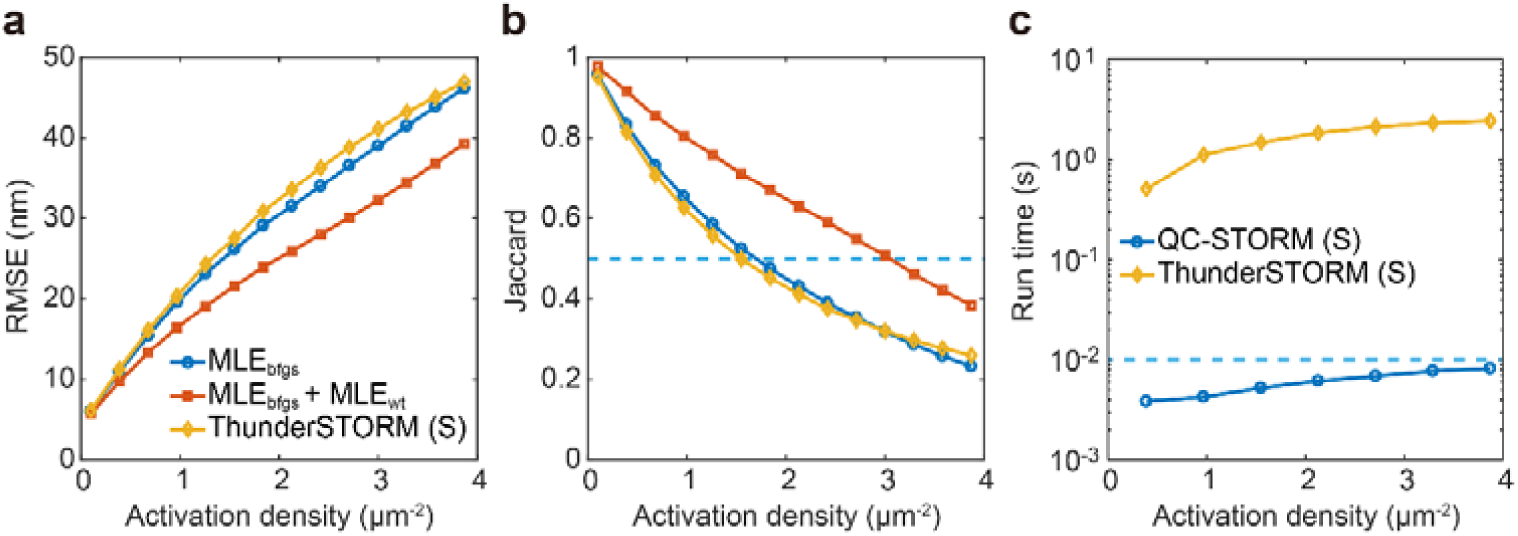
Comparison on the localization performance among MLE_bfgs_, MLE_wt_ and ThunderSTORM for simulated 2D raw images. (a) The dependance of RMSE on activation density. (b) The dependance of Jaccard index on activation density. (c) The dependance of run time on activation density. Here the run time accounts for all data processing routine, including ROI extraction, molecule localization, and super-resolution image rendering, to process a single raw image. The simulated images contains 1024 × 1024 pixels with a pixel size of 100 nm. Both the QC-STORM and the ThunderSTORM were set to work at the sparse emitter mode when the run time was evaluated. Both the MLE_bfgs_ and the MLE_wt_ algorhthms were used when the QC-STORM is working in the sparse emitter mode. The letter S denotes sparse emitter mode.

**Fig. 5.**
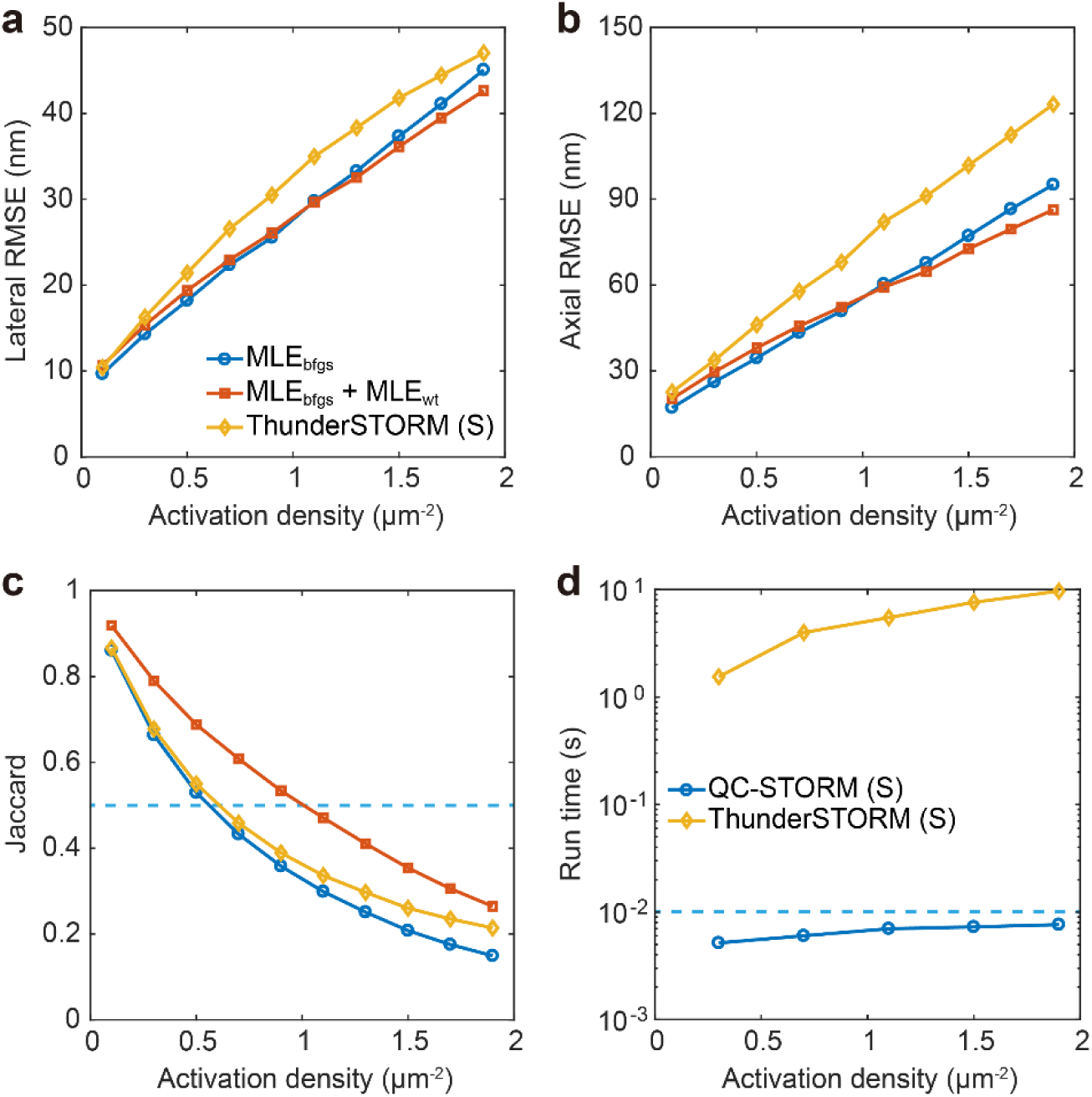
Comparison on the localization performance among MLE_bfgs_, MLE_wt_ and ThunderSTORM for simulated astigmatism 3D raw images. (a) The dependance of lateral RMSE on activation density. (b) The dependance of axial RMSE on activation density. (c) The dependance of Jaccard index on activation density. (d) The dependance of run time on activation density. Here the run time accounts for all data processing routine in a single raw image. Other settings are similar to those in Fig. 4.

It is interesting to evaluate the data processing speed of our MLE-based algorithms. For 2D imaging where the ROI size is 7 × 7 pixels, we found that the localization speed is 8.7 × 10^6^ ROIs per second for MLE_bfgs_ and 8.2 × 10^6^ ROIs per second for MLE_wt_, respectively. Compared with MATLAB implemented MLE_bfgs_, which only achieves a localization speed of 150 ROIs per second, the speed is increased by 5.8 × 10^3^ times after applying GPU computation. Meanwhile, for astigmatism 3D imaging where the ROI size is 11 × 11 pixels, we achieve a localization speed of 2.4 × 10^6^ ROIs per second when MLE_wt_ is used.

We further compared the full data processing routine, rather than the localization speed only, between our QC-STORM and the popular ThunderSTORM, using simulated raw images with a size of 1024 × 1024 pixels (corresponding to an FOV of 102 μm × 102 μm, when the pixel size is 100 nm). We found that, when both plug-ins are working at the sparse emitter mode, QC-STORM is about two orders of magnitude faster than ThunderSTORM for both 2D and astigmatism 3D raw images (Fig. 4c & Fig. 5d). These data also point out that QC-STORM is capable of achieving real-time processing within 10 ms exposure time for an activation density of up to 3 molecules / μm^2^ for 2D imaging (Figs. 4b-4c), and 1 molecules / μm^2^ for astigmatism 3D imaging (Figs. 5c-5d), while the detection rate is still acceptable (that is, the Jaccard index is better than 0.5). Interestingly, although Daniel et al mentioned that the reported algorithms for 2D localization of sparse emitters is near optimal [29], here we show an improved localization precision and/or detection rate after combining weighted MLE (MLE_wt_) with conventional MLE (MLE_bfgs_), with only a slight speed degradation when using weighted MLE.

### 3.3 Localization of multiple emitters using simulated datasets

As seen in Fig. 3a, we combined the divide and conquer strategy with a series of MLE-based localization algorithms to process raw images with high activation density. Therefore, it would be more useful to assess the overall performance of QC-STORM working at multi-emitter mode (corresponding to a combination of MLE_bfgs_, MLE_wt_, MLE_2e_ and MLE_3e_), rather than individual multi-emitter algorithms (that is, MLE_2e_, MLE_3e_). Using simulated 2D and astigmatic 3D raw images, we compared the localization performance of QC-STORM with ThunderSTORM and found that ThunderSTORM exhibits slightly better performance in both localization precision (Fig. 6a, Fig. 7a-7b) and detection rate (Fig. 6b, Fig. 7c), but runs at three orders of magnitude slower speed than QC-STORM (Fig. 6c, Fig. 7d). For the comparison between QC-STORM and WindSTORM using 2D simulated images, WindSTORM presents better localization precision and detection rate than QC-STORM (Figs. 6a-6b).

**Fig. 6.**
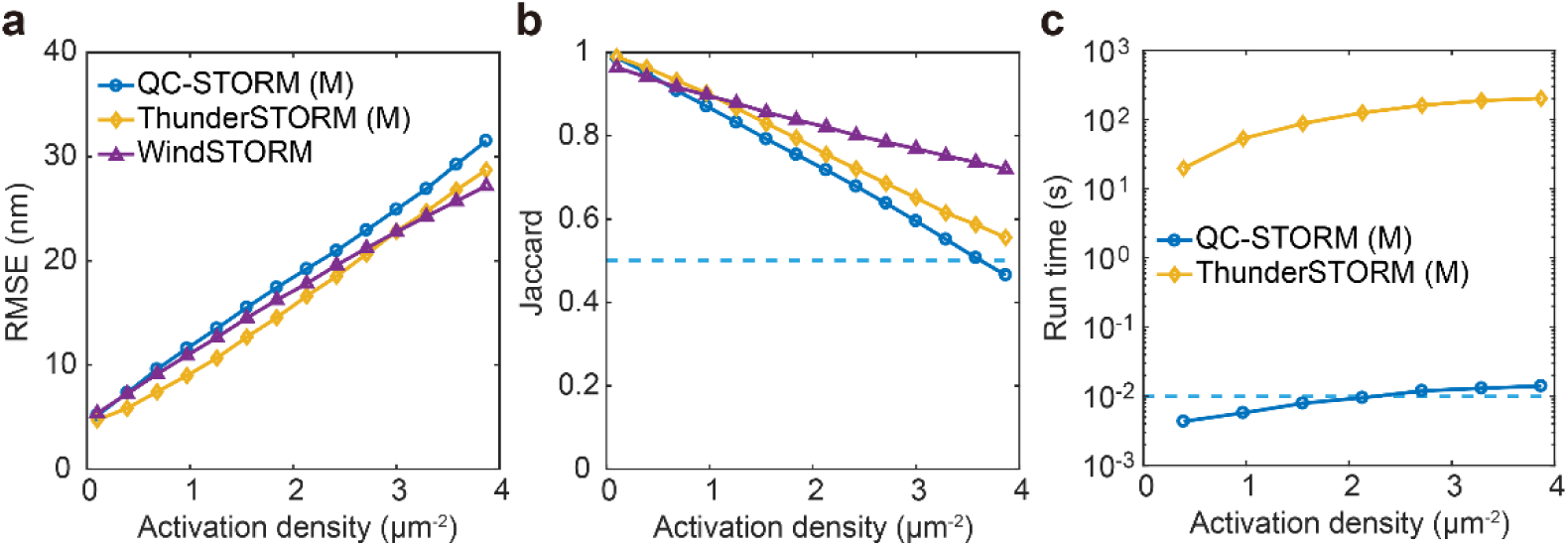
Comparison on the localization performance of three software for 2D raw images. (a) The dependance of RMSE on activation density. (b) The dependance of Jaccard index on activation density. (c) The dependance of run time on activation density. Here the run time accounts for all data processing routine in a single raw image. Other settings are similar to those in Fig. 4. The letter M denotes multi-emitter mode, and WindSTORM has only a multi-emitter mode. We don’t show the run time of WindSTORM, since the GPU version of WindSTORM is not running correctly under our hardware configurations.

**Fig. 7.**
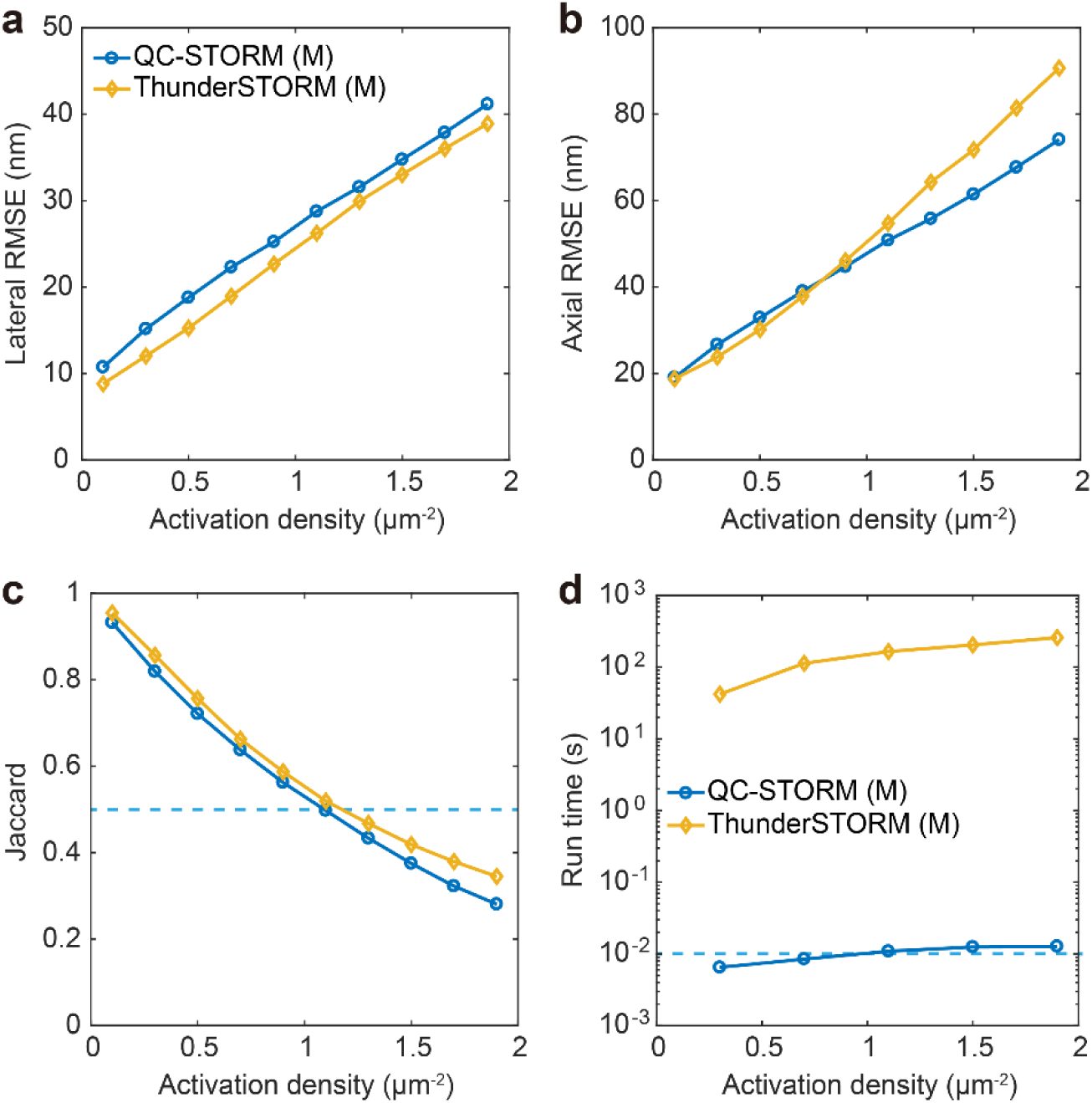
Comparison on the localization performance of ThunderSTORM and QC-STORM for astigmatism 3D raw images. (a-b) The dependance of lateral and axial RMSE on activation density. (c) The dependance of Jaccard index on activation density. (d) The dependance of run time on activation density. Here the run time accounts for all data processing time in a single image of 1024 × 1024 pixels. The letter M denotes multi-emitter mode.

Because the current GPU version of WindSTORM is not compatible with our Dell precision T7610 workstation and GeForce RTX 2080 Ti graphic card, we estimated the run time of WindSTORM using the previously reported value, after considering the difference of image size. The run time of the GPU-based WindSTORM was previously measured with a GTX 1080 graphics card, and could possibly have a 50% reduction when a RTX 2080 Ti graphics card is used. Therefore, although the run time of WindSTORM may not depend linearly on the performance (or single floating-point operation speed) of GPU, we can still roughly estimate that WindSTORM runs slightly slower than QC-STORM, after using GPU to acceralate both software. Note that, in this study the localization precision and detection rate of WindSTORM were measured using its MATLAB version, and that currently WindSTORM is not capable of processing 3D images.

Considering that QC-STORM provides advantageous performance in the run time, we further investigated the localization speed of our MLE-based localization algorithms for two emitters (MLE_2e_) and three emitters (MLE_3e_). For 2D imaging with an ROI size of 7 × 7 pixels, we achieve a localization speed of 3.7 × 10^6^ and 1.9 × 10^6^ ROIs per second for MLE_2e_ and MLE_3e_, respectively. And, for astigmatism 3D imaging with an ROI size of 11 × 11 pixels, we achieve a localization speed of 7.6 × 10^5^ ROIs per second for two-emitter localization. These findings indicate that our MLE-based algorithms for multiple emitter run at only 2-4 times slower speed than the fast MLE-based algorithms for sparse emitters (that is, MLE_bfgs_ and MLE_wt_).

We also compared the localization performance of QC-STORM with ThunderSTORM and 3D-DAOSTORM using open datasets [29]. The open datasets contain 2D and astigmatism 3D images at both low activation density and high activation density, and low SNR and high SNR. The results are shown in Fig. 8. For both 2D and astigmatism 3D images, the localization precision and detection rate of QC-STORM are comparable to those of ThunderSTORM, and are not as good as 3D-DAOSTORM [33]. It should be noted that the reported localization performance of QC-STORM [29] was tested using only MLE_bfgs_ under a relative low-end graphics card (compared with RTX 2080 Ti in this study). The updated results have been recently uploaded to the website using the name of QC-STORM-2019 [34].

**Fig. 8.**
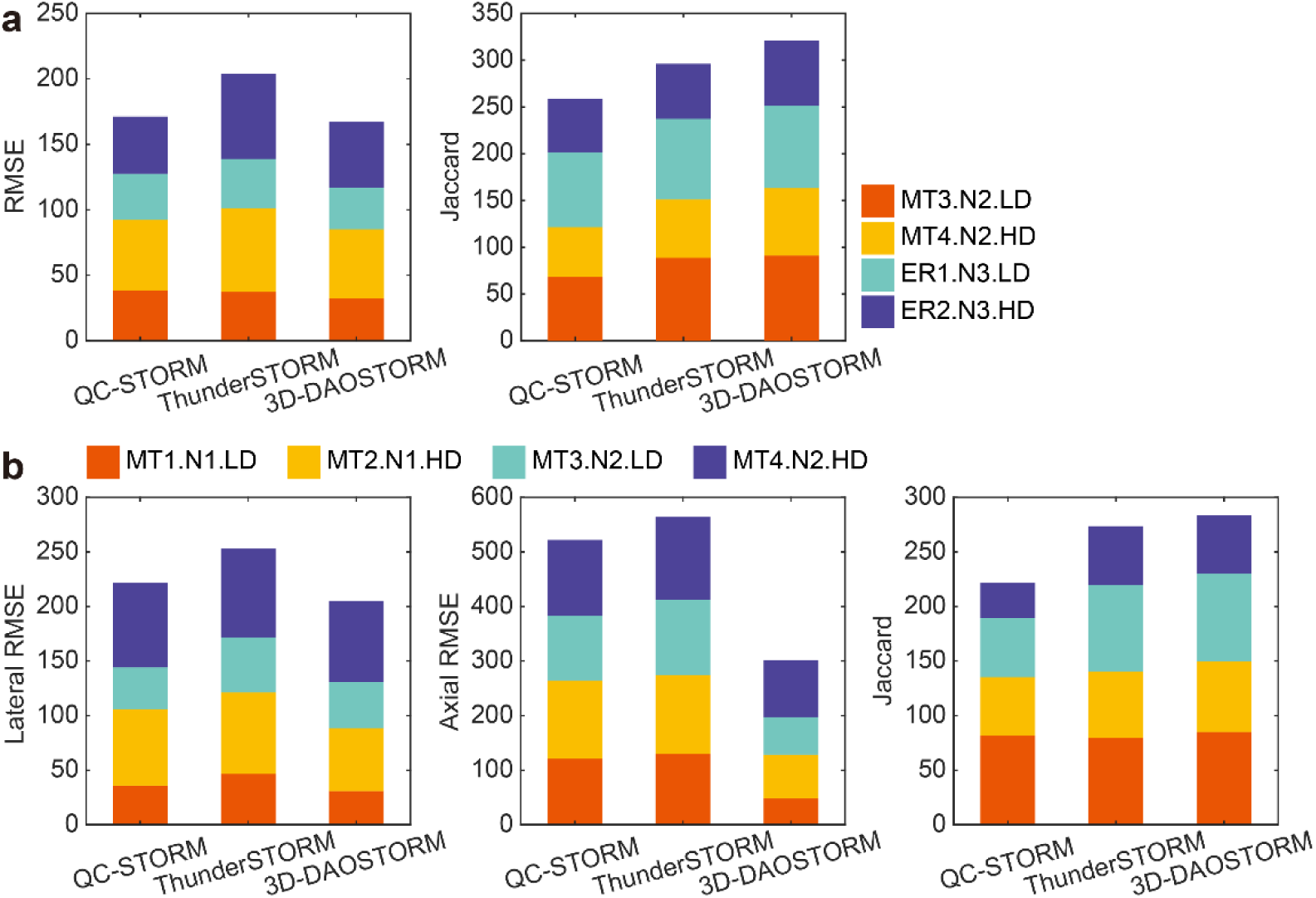
Comparison on the localization performance of three software using open datasets. (a) Evaluation of RMSE and Jaccard index on four 2D imaging datasets. (b) Evaluation of lateral and axial RMSE, and Jaccard index on four astigmatism 3D imaging datasets. The acronym LD and HD denote low density and high density, respectively. MT1 and MT2 are high SNR datasets. MT3, MT4, ER1 and ER2 are low SNR datasets.

For the computation speed, we failed to run 3D-DAOSTORM using our simulated data. However, 3D-DAOSTORM has been shown to have 2-3 orders of magnitude slower speed than the GPU accelerated WindSTORM [6], while the latter runs at similar speed as our QC-STORM.

### 3.4 Testing QC-STORM using experimental data

We performed 2D STORM imaging of the microtubules in U-2 OS cells, and captured a total of 1000 raw images with an image size of 1024 × 1024 pixels and a pixel size of 108.3 nm (corresponding to an FOV of 110 μm × 110 μm) under high activation density (Fig. 9a). We found that QC-STORM is able to localize a total of 6.6 × 10^3^ molecules per raw image, and spends 6 seconds to finish the full processing of 1000 raw images (including ROI extraction, molecule localization and super-resolution image reconstruction). Therefore, the total image processing time per raw image is 6 ms, which is less than the exposure time of 10 ms. For WindSTORM, the reported time for processing 2000 images with an image size of 512 × 512 pixels is 6.5 seconds [6], corresponding to 13 seconds for processing 1000 images with an image size of 1024 × 1024 pixels. This time can be roughly shorten to 6.5 seconds after replacing the previous GTX 1080 graphics card with the current RTX 2080 Ti graphics card. For ThunderSTORM, we estimated that a total of 22 hours is required to process 1000 raw images with 1024 × 1024 pixels in each image. Therefore, the full processing speed of QC-STORM is 1.32 × 10^4^ times over ThunderSTORM, and is slightly faster than WindSTORM. The total image processing times for the three software are shown in Fig. 9a.

**Fig. 9.**
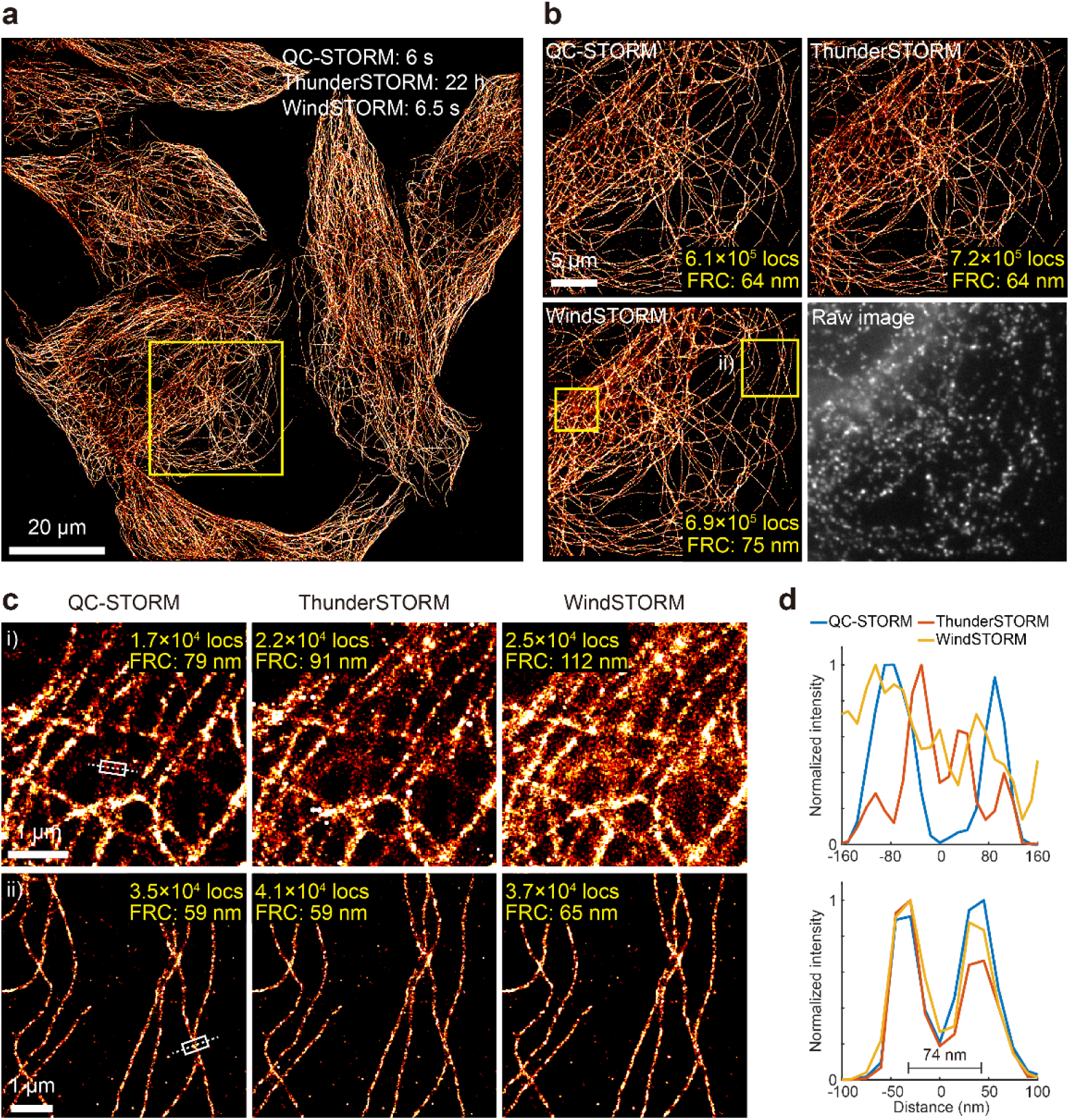
Evaluation of the localization performance among QC-STORM, ThunderSTORM and WindSTORM using experimental 2D images. (a) A super-resolution image reconstructed from a total of 1000 raw images with 1024 × 1024 pixels. The images were processed by QC-STORM. The estimated full data processing times for different software are shown in the top-right corner. (b) Enlarged super-resolution images of the boxed region in (a). A representative raw image shows a non-uniform fluorescence background. Three software were used to process the raw images independently. The FRC resolution and the total number of localizations are shown in the lower-right corners. c) Enlarged super-resolution images of the two ROIs indicated in (b). The upper row shows an area with high fluorescence background, and the lower row is for an area with low fluorescence background. The localization number and the FRC resolution of these ROIs are also shown in these images. (d) Spatial resolution characterized by the intensity profiles of the marked positions in (c).

We calculated the total number of localized molecules and the FRC resolution of a zoom-in image with 256 × 256 pixels (Fig. 9b). We found that QC-STORM localizes a total of 6.1 × 10^5^ molecules, which is less than that from ThunderSTORM (7.2 × 10^5^) or WindSTORM (6.9 × 10^5^). However, the FRC resolution of QC-STORM (64 nm) is the same as that from ThunderSTORM (64 nm), but is significantly better than that from WindSTORM (75 nm). Note that WindSTORM achieves a slightly higher localization precision than QC-STORM in simulated data, but a lower FRC resolution in this experiment data. This is probably due to: 1) the SNR of the experimental data is much more complex than that in the simulated data, and thus results in localizations with different localization precision; and 2) QC-STORM filters out localizations with low precision (see the source codes of QC-STORM for more details: https://github.com/SRMLabHUST/QC-STORM.). Actually, for an ROI with high fluorescence background (Fig. 9c), which usually suffers from low localization precision, the super-resolution image reconstructed from QC-STORM shows fewer scattered localizations than those from ThunderSTORM and WindSTORM. As a result, the FRC resolution achieved by QC-STORM is higher than those from both ThunderSTORM and WindSTORM (Fig. 9c). On the other hand, for an ROI with low fluorescence background (Fig. 9c), QC-STORM exhibits the same FRC resolution as ThunderSTORM, and slightly higher FRC resolution than WindSTORM. These results are also verified by the intensity profile analysis of two cross-linked microtubules (Fig. 9d).

We also evaluated the localization performance of QC-STORM using an experimental open dataset with sparse emitters from astigmatism 3D imaging (Fig. 10a) [29]. QC-STORM requires 34 s to process the 112683 images with 324 × 271 pixels in each image, corresponding to a full data processing time of 0.3 ms per image. For the same open dataset, ThunderSTORM requires 4.6 hours to finish the full image procession, showing a much slower time of 146 ms per image. Moreover, QC-STORM achieves an FRC resolution of 36 nm, which is better than that from ThunderSTORM (44 nm). The better resolution of QC-STORM over ThunderSTORM is also verified by the line profile analysis of microtubules (Fig. 10b-10c, upper row), where the two microtubules are clearly resolved from QC-STORM, but are blurred from ThunderSTORM. The reason is that MLE_wt_ in the QC-STORM plug-in provides more reliable localizations for contaminated molecules, which occur with high probability at the cross-linked region of microtubules. Additionally, we observe that the calculated depths of the microtubules are slightly different between QC-STORM and ThunderSTORM (Fig. 10b). This difference probably results from the different methods for calibration curve measurement. The calibration curve in our QC-STORM is measured between z depth and 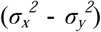 [35], while ThunderSTORM adopt the calculation method from Huang et al [19]. Actually, the same open dataset has been analyzed by several software [34], and the reported depths are different. However, it is hard to determine which result is more accurate, since no ground-truth is provided for this experimental open dataset.

**Fig. 10.**
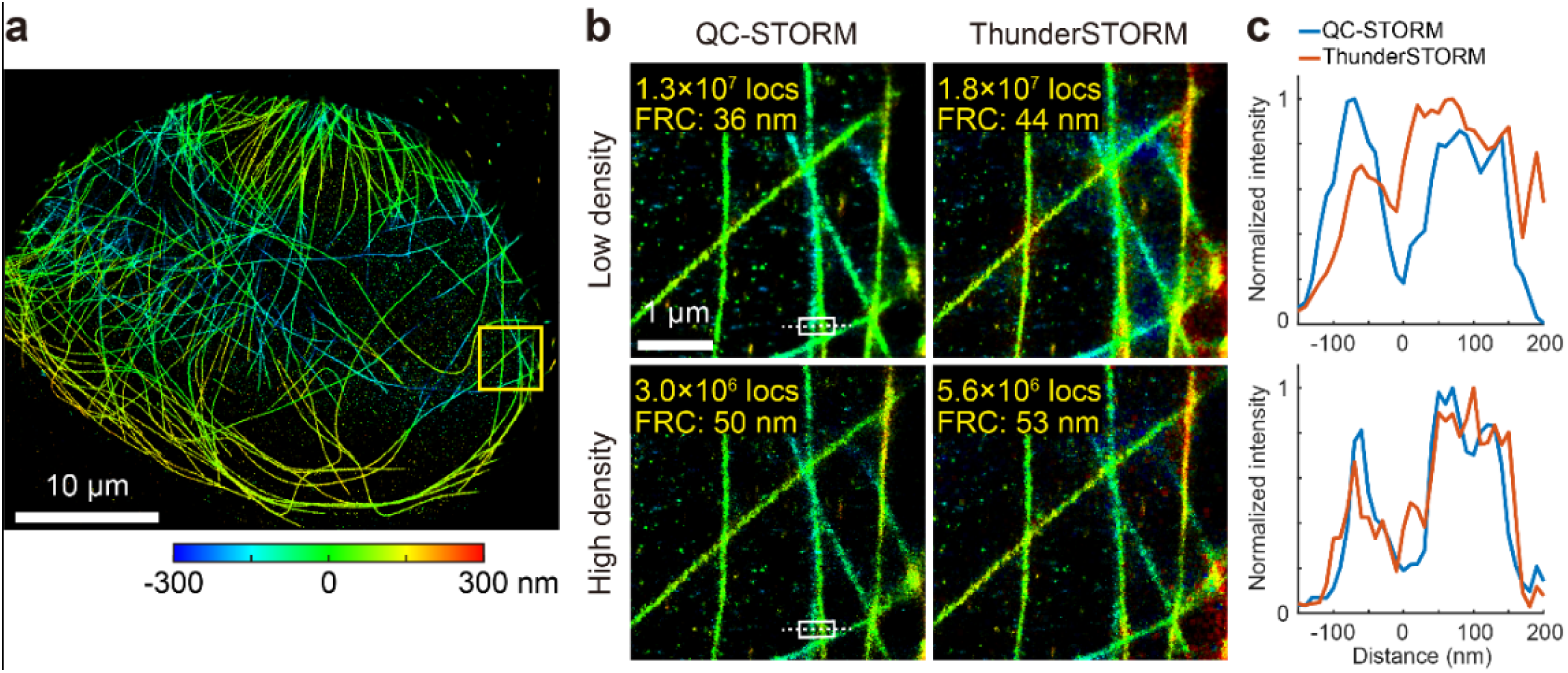
Evaluation of the localization performance between QC-STORM and ThunderSTORM using an experimental 3D open dataset. (a) A super-resolution image recontructed from QC-STORM. (b) Enlarged super-resolution images reconstructed from the two software. The images correspond to the marked area in (a). The upper images were processed from the original open dataset with low activation density, while the lower images were processed from a modified open dataset with high activation density. The FRC resolution and the total localization number are shown in the upper-left corners. (c) Spatial resolution characterized by the intensity profiles of two microtubules at the marked positions in (b).

To evaluate the performance of QC-STORM in processing 3D raw images with high activation density, we modified the experimental open dataset by merging every 10 images into one image and obtain 11268 raw images. We found that QC-STORM takes 8 s to process the modified dataset (or 0.7 ms per image), while ThunderSTORM spends 26 hours for the same dataset (or 8 s per image). Moreover, the FRC resolution achieved by QC-STORM is 50 nm, which is slightly higher than that by ThunderSTORM (53 nm). The intensity profile analysis on two closely microtubules also confirms this finding (Fig. 10b-10c).

### 3.5 QC-STORM for real-time image acquisition feedback

Benefiting from the fast data processing speed, QC-STORM can provide real-time statistics of many parameters (for example, fluorescence signal intensity, photon background intensity, SNR, PSF width, localization precision). These statistics are useful for enabling online data acquisition optimization [11-13]. Here we demonstrate that SNR can be used for active axial drift correction, and the number of localized molecules can be used for optimizing activation laser intensity.

The axial drift correction is based on real-time maximization of SNR during image acquisition. Compared with the drift correction method using fiducial markers or bright field images [36-38], our method doesn’t require extra costs or efforts in the sample preparation or imaging system modification, thus is beneficial for cost-efficient localization microscopy [39, 40]. Our drift correction method is described below. After acquiring every 500 images (or other user-defined number, depending on the exposure time for each raw image and the stability of the imaging system), QC-STORM drives a step motor (SS1704A20A with MD-2322 controller, Samsr motor, Japan) which is connected to the focusing knob of the microscope. Raw images are captured from 5 successive focal planes after moving the focusing knob with 30 nm step size, and then analyzed using QC-STORM to find a focal plane where the SNR is maximal. This process can be repeated several times for correcting larger focal drift or moving to a new FOV. Moreover, the drift correction is made to be automatic and the raw images used for the drift correction are not mixed with the raw images for SRLM experiments. That is to say, the axial drift correction from QC-STORM is almost transparent to SRLM experiments. We assessed the validity of this axial drift correction method by comparing 2D imaging results with or without axial drift correction. A total of 8000 raw images were acquired and analyzed from the microtubules in U-2 OS cells. The results show that both the SNR and the total photon number are kept stable during the whole image acquisition process when the axial drift correction is applied, while these two parameters decrease notably (15% – 30%) when the axial drift correction is turned off (Fig. 11a-11b).

**Fig. 11.**
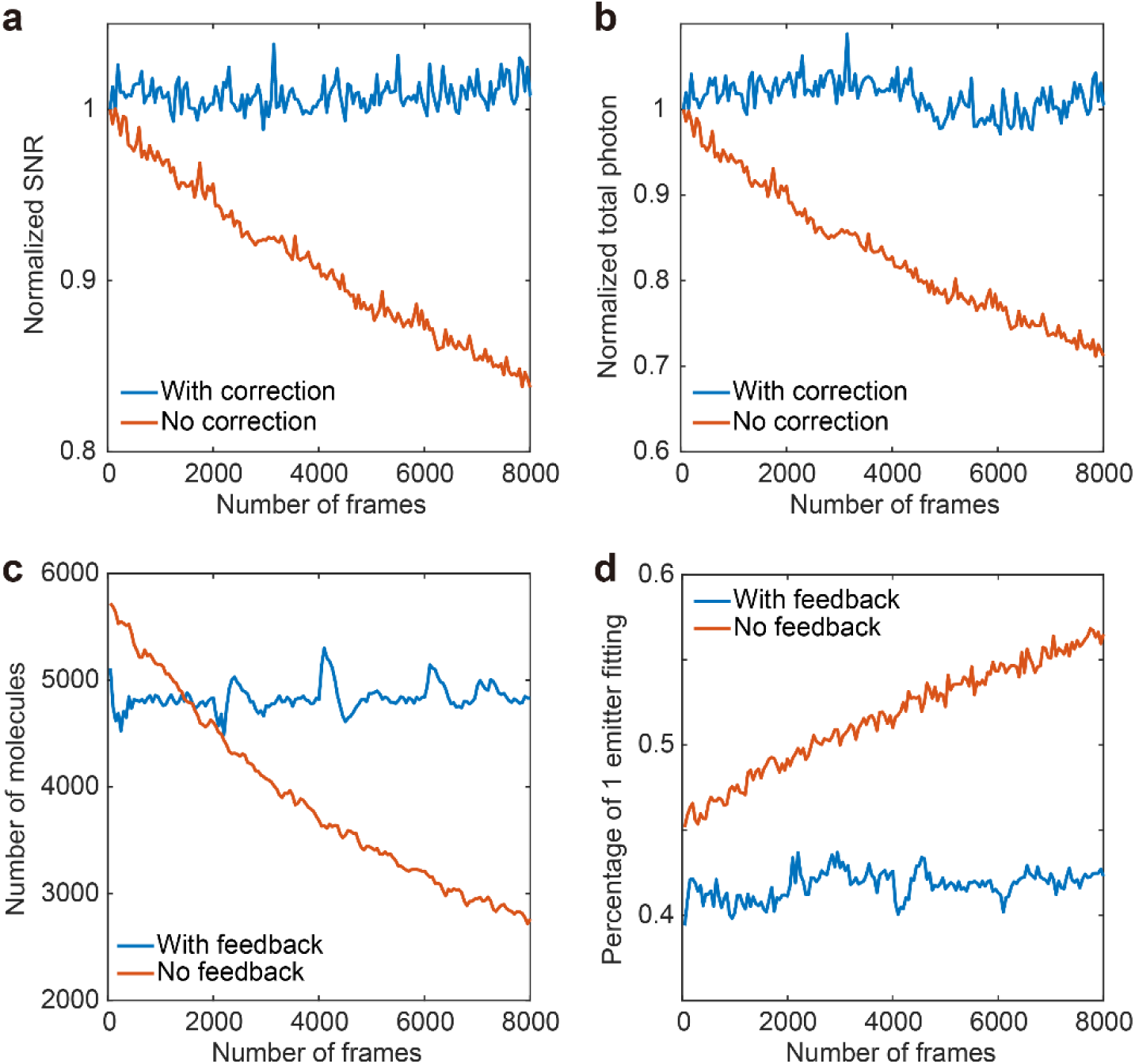
Real-time feedback control on image SNR and activation density. (a-b) The dependence of normalized SNR and total signal photon on the number of raw image frames. The axial drift correction is turned ON (with correction) or OFF (no correction). (c-d) The dependence of the number of localized molecules and the percentage of sparse emitter localization on the number of raw image frames. The activation density control is turned ON (with feedback) or OFF (no feedback).

We can also use QC-STORM to provide feedback for controlling the intensity of activation laser, thus stabilizing the activation density of molecules. Firstly, we estimate the activation density using the percentage of single emitter localization (see Fig. 3b). Then, we calculate the error between the observed activation density and the user expected activation density. The error is used as the input in a proportional–integral–derivative (PID) controller, which is further used to adjust the activation laser intensity via an analog-to-digital converter. For simplicity, the derivative coefficient is set to 0, and the proportional and integral coefficients are acquired by preliminary experiments to enable a stable and quick response. Through experimental 2D imaging of the microtubules in U-2 OS cells, we verify that the number of activated molecules (Fig. 11c) and the percentage of single molecule localization (Fig. 11d) in each raw image can be both stabilized during the whole image acquisition process, after enabling the activation density feedback. In contrast, when the activation density feedback is not applied, the number of activated molecules per frame decreases by 50% at the end of the experiment (Fig. 11c). Note that the jump in the number of activated molecules is due to overcompensation from the axial drift correction.

### 3.6 Combing QC-STORM with ANNA-PALM for higher imaging speed

QC-STORM enables real-time image processing for ROIs containing multiple emitters, thus significantly improves the imaging speed of SRLM for a given resolution [23, 41]. On the other hand, a deep-learning based method called ANNA-PALM is able to generate a super-resolution image using a significantly smaller number of localizations, thus massively increasing the imaging speed compared with traditional methods using sparse emitter localization. Considering that ANNA-PALM still requires a sufficient number of localizations from conventional localization algorithms, here we try to combine QC-STORM with ANNA-PALM to provide a further improvement in the imaging speed. Using images with an activation density of 3.5 × 10^3^ molecules per image (Fig. 12), we investigated the relationship between the quality of reconstructed super-resolution images and the number of raw images. We analyzed the intensity profile analysis of two cross-linked microtubules (Fig. 12a-12b), and found out that QC-STORM requires a total of 800 images to resolve the two microtubules, while a combination of ANNA-PALM with QC-STORM can resolve the structure using only 300 images (Fig. 12c). However, when combining ANNA-PALM with sparse emitter localization (as demonstrated in the original paper [10]), a total of 1000 raw images would be required to reconstruct a super-resolution image with similar resolving power. That is to say, the imaging speed of ANNA-PALM could be further improved (3 times in this study) after combining ANNA-PALM with a suitable multi-emitter localization algorithm (QC-STORM in this study). And, it is worthy to point out that the combined approach provides a larger separation distance for the two microtubules (Fig. 12c, 105 nm from QC-STORM and 118 nm from the combination), indicating a further need to investigate the benefits and drawbacks from the combined approach. Moreover, ANNA-PALM requires 5 minutes to process the super-resolution image shown in Fig. 12a (80 μm × 86 μm, 20 nm pixel size), which is apparently too slow to be used in real-time image acquisition optimization during SRLM experiments.

**Fig. 12.**
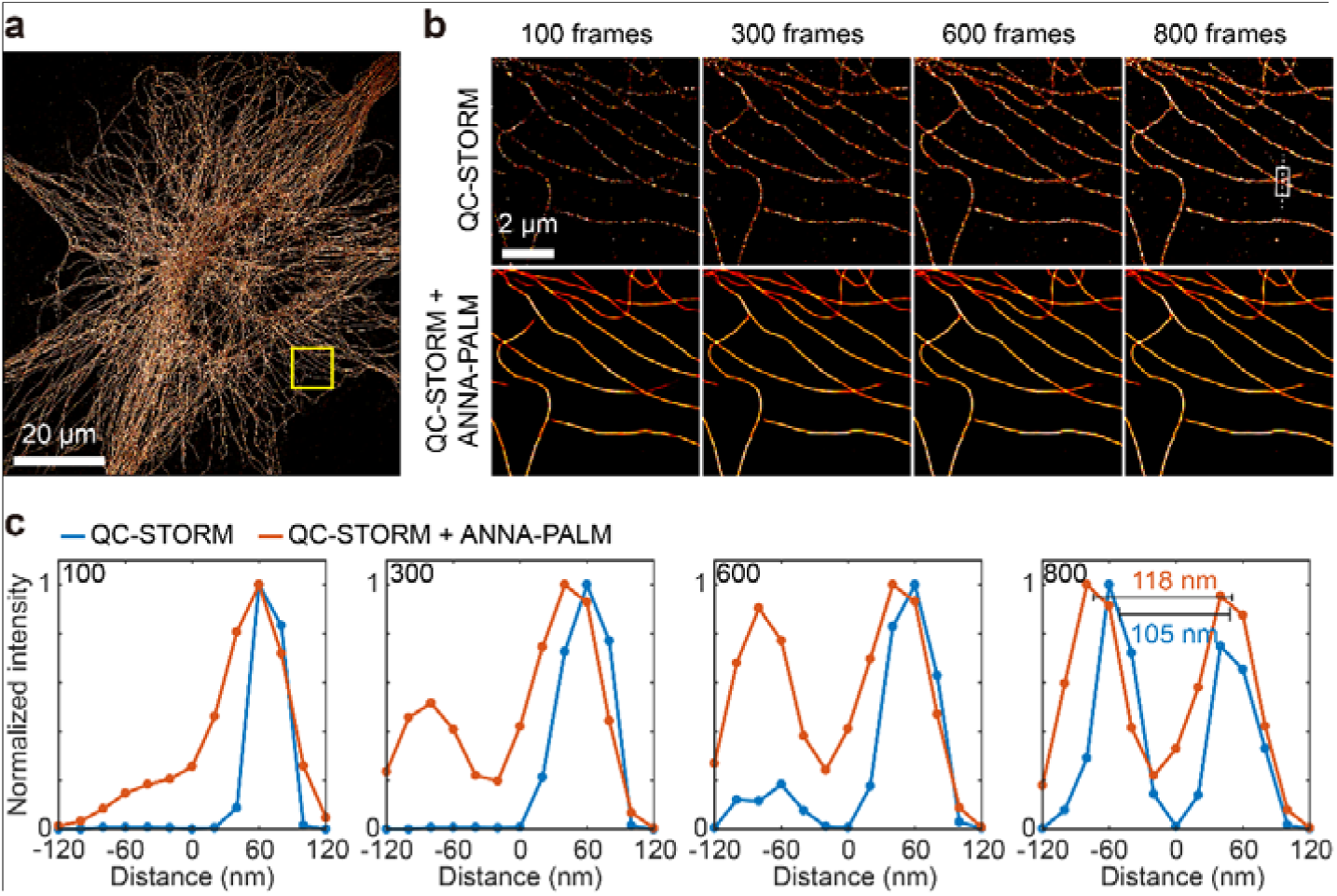
The improvement of imaging speed after combining ANNA-PALM with QC-STORM. (a) A super-resolution image reconstructed from QC-STORM. Here a total of 800 raw images was used, and the FOV equals to 80 μm × 86 μm. (b) Super-resolution images reconstructed from different number of raw images. These images were originated from the rectangular area marked in (a). QC-STORM or a combination of QC-STORM and ANNA-PALM was used to process the raw images. (c) Line profiles of the marked areas in (b).

## 4. Conclusion

We adopt the divide and conquer strategy and present a user-friendly Java-based plug-in called QC-STORM that is capable of providing fast MLE fitting of multiple emitters. QC-STORM is consisted of a series of MLE-based localization algorithms, including MLE_bfgs_, MLE_wt_, MLE_2e_ and MLE_3e_, and is capable of processing simulated and experimental 2D and 3D ROIs with 3-4 orders of magnitude faster speed than the popular fitting-based ThunderSTORM, with only a slightly worse localization precision. Compared with the newly reported non-iterative WindSTORM, QC-STORM exhibits a similar image processing speed, with a reduction on localization precision and detection rate. Note that WindSTORM for 3D imaging is not currently available, probably because the complicated pre-processing procedures in the ROI identification results in distorted emission patterns for 3D localization.

We demonstrate that QC-STORM could provide real-time feedback for active axial drift correction and activation density optimization. We also show that combining QC-STORM with ANNA-PALM could further increase the imaging speed. We provide user-friendly plugins of QC-STORM for ImageJ and Micro-Manager software.

Finally, we point out that the performance of the current QC-STORM is still limited by the setting of maximum number of emitters to be localized (that is, 3 emitters for 2D imaging and 2 emitters for 3D imaging), because currently we aim to provide real-time data processing and image acquisition optimization for raw images with 1024 × 1024 pixels and 10 ms exposure time (which is popularly used in SRLM with sCMOS cameras). Further improvement on the performance of QC-STORM could be made possible by using multiple GPU, or a better pre-processing method for ROI identification.

## Acknowledgments

We thank Daniel Sage for evaluating QC-STORM with open datasets.

